# Serverless Prediction of Peptide Properties with Recurrent Neural Networks

**DOI:** 10.1101/2022.05.18.492545

**Authors:** Mehrad Ansari, Andrew D. White

## Abstract

We present three deep learning sequence prediction models for hemolysis, solubility, and resistance to non-specific interactions of peptides that achieve comparable results to the state-of-the-art models. Our sequence-based solubility predictor, MahLooL, outperforms the current state-of-art methods for short peptides. These models are implemented as a static website without the use of a dedicated server or cloud computing. Web-based models like this allow for accessible and effective reproducibility. Most existing approaches rely on third-party servers typically that require upkeep and maintenance. That trend leads to a relatively longer lifetime of web-based models. These predictive models do not require servers, require no installation of dependencies, and work on across a range of devices. The models are bidirectional recurrent neural networks. This *serverless* prediction model is a demonstration of edge machine learning that removes the dependence on cloud providers. The code and models are accessible at https://github.com/ur-whitelab/peptide-dashboard.

## 1 Introduction

Deep learning models have been widely applied to extract information from big data in cheminformatics. Compared to machine learning algorithms, deep learning can perform feature extraction and learn patterns over various nonlinear layers of representations of the input data,^[1]^ can explain the vanishing effects of gradients,^[2]^ and perform better with raw high-dimensional data.^[3]^ There is a growing increase in the number of web-based implementations of deep learning frameworks that provide convenient public access and ease of use.^[4–9]^ Notably, many web servers have been developed for sequence design tasks, like analysis of RNA, DNA, or proteins. For example, survival analysis based on mRNA data (GENT2,^[10]^ PROGgeneV2,^[11]^ SurvExpress,^[12]^ MEXPRESS,^[13]^ etc.), studying prognostic implications of non-coding RNA (PROGmiR,^[14]^ SurvMicro,^[15]^ OncoLnc,^[16]^ TANRIC^[17]^), survival analysis based on protein (TCPAv3.0,^[18]^ TRGAted^[19]^) and DNA (MethSurv,^[20]^ cBioPortal^[21]^) data, and multiple areas of assessing cancer therapeutics.^[22]^ These scientific web servers and web-based services allow for the availability of complex inference algorithms to a much broader user community and promote open science. This is especially important because of the disparities between lower and higher income nations, where there are disparities in the types of research activities that can be performed.^[23]^ Cheminformatics-related research, the topic of this work, mostly takes place at those nations privileged with resource-rich institutions, where there are adequate funding resources. Yet, web-based implementations can broaden access to these methods.

Beyond disparities among institutions, web-based implementations are also a mechanism for reproducibility in science. In peptides specifically, Melo et al.^[24]^ argue that deep learning sequence design should be accomplished by free public access to the (1) source code, (2) training and testing data, (3) published findings. However, this is not often true; Littmann ^[25]^ found in an analysis of ML research articles in bio-medicine and life sciences published between 2011 and 2016 that only 50% released software, while 64% released data. Web-based servers do not fit the exact definition of open science (due to lack of source code access), but they do accomplish the goal of enabling others with broader expertise to build on previous advances, and are often more accessible and convenient than access to model and source code alone.

Thus, there is a compelling argument to continue web-based tools. There are, however, two major drawbacks: source-code can be inaccessible as discussed above and the reliance on third-party or self-hosted servers. Deep learning inference often requires GPUs, and this requires a specialized hosting service or a complex self-hosted set-up. This creates difficult ongoing expenses, and many tools are thus only available for a limited time after publication. Additionally, there can be low incentives to increase capacity. Popular tools, like RoseTTAFold,^[26]^ can have days-long queues. The expense and deployment problems also can create disparities in impact of research between resource-rich and low-resource institutions, because not all researchers can afford to create web-based implementations.

To address the challenges above, we demonstrate a *serverless* deep learning web-based server, https://peptide.bio, that predicts peptide properties using recurrent neural networks (RNN) via users’ local devices. These trained models are implemented in JavaScript and are loaded to a user’s web browser. Users make predictions by running these trained models on a web browser on their local machines, or even cell phones, without having to install any modules. They can be run locally as well, if desired^1^ The *serverless* computing describes a programming model and architecture, where small code snippets are executed in the cloud without any control over the resources on which the code runs.^[27]^ This is by no means an indication that there are no servers. Simply, it means that the developer leaves most operational concerns such as resource provisioning and scalability to the cloud provider (the end-user in our case). John et al.^[28]^ proposed the idea of serverless computing for efficient model utilization on could resources without specific constraints on the cloud provider. However, in this work, we seek to fully remove the need for a cloud provider, and bypass this conventional dependency. While this is indeed impractical for the resource-intensive training step of deep learning models, the trained models are typically cheap to evaluate, thus, inference is robustly feasible on even limited computing resources (i.e. commodity phones, laptops). This removes hosting costs and the conventional dependence on cloud providers or self-hosting of resource-rich academic institutions. Although we make some compromises here on model size and complexity, we expect the continued improvement of hardware (i.e. Moore’s law^[29]^) to increase the type of models possible in JavaScript each year. This serverless approach should accelerate reproducible ML science, while also lowering the gap between resource-rich universities and the rest, as well as enabling a better dissemination of research from a broader community of chemists.

This manuscript is organized as follows: We start by providing a brief overview of some comparable predictive sequence-based models for the classification tasks in this work (hemolysis, solubility and non-fouling) in Section 1.1. In Section 2, we describe the datasets, architecture of our deep learning models, the choices for the hyperparameters, as well as a high level overview of the methods used in the previous comparable sequence-based models in the literature. This is followed by evaluating the model in a comparative setting with the state-of-art models in Section 3. Finally, we conclude the paper in Section 4, with a discussion of the implications of our findings.

### 1.1 Previous Work

Quantitative structure–activity relationship (QSAR) modelling is a well-established field of research that aims at mapping sequence and structural properties of chemical compounds to their biological activities.^[30]^ QSAR models have been successfully applied to angiotensin converting enzyme(ACE)-inhibitory peptides,^[31–33]^ antimicrobial peptides,^[34–37]^ and antioxidant peptides.^[38–40]^ For solubility predictions, DSResSol (1)^[41]^ improved prediction accuracy (ACC) and AUROC to 75.1% and 0.84, respectively, by identifying long-range interaction information between amino acid k-mers with dilated convolutional neural networks and outperformed all existing models such as DeepSol,^[42]^ PaRSnIP,^[43]^ SoluProt,^[44]^ Protein–Sol^[45]^ and PROSO II.^[46]^ HAPPENN^[47]^ forms the state-of-art model for hemolytic activity prediction with ACC of 85.7% and has better performance compared with HemoPI^[48]^ and HemoPred.^[49]^ Hasan et al.^[50]^ developed a two-layer prediction framework, called HLPpred-Fuse, that can distinguish between hemolytic and non-hemolytic peptides, as well as their low and high activity, with an AUROC of 0.91 (averaged over two reported independent datasets). However, short peptides (*<*6 amino acid residues) are excluded from their datasets due to the difficulty in capturing meaningful sequence information from shorter peptides.

## 2 Materials and Methods

### 2.1 Datasets

#### 2.1.1 Hemolysis

Hemolysis is defined as the disruption of erythrocyte membranes that decrease the life span of red blood cells and causes the release of Hemoglobin. Identifying non-hemolytic antimicrobial is critical to their applications as non-toxic and safe measurements against bacterial infections. However, distinguishing between hemolytic and non-hemolytic peptides is complicated, as they primarily exert their activity at the charged surface of the bacterial plasma membrane. Timmons and Hewage^[47]^ differentiate between the two whether they are active at the zwitterionic eukaryotic membrane, as well as the anionic prokaryotic membrane. In this work, the model for hemolytic prediction is trained using data from the Database of Antimicrobial Activity and Structure of Peptides (DBAASP v3^[51]^). The activity is defined by extrapolating a measurement assuming dose response curves to the point at which 50% of red blood cells (RBC) are lysed. If the activity is below 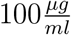, it is considered hemolytic. Each measurement is treated independently, so sequences can appear multiple times. The training data contains 9,316 sequences (19.6% positives and 80.4% negatives) of only L- and canonical amino acids. Note that due to the inherent noise in the experimental datasets used, in some observations (*∼*40%), an identical sequence appears in both negative and positive class. As an example, sequence “RVKRVWPLVIRTVIAGYNLYRAIKKK”, is found to be both hemolytic and non-hemolytic in two different lab experiments (i.e. two training examples).

#### 2.1.2 Solubility

The training data contains 18,453 sequences (47.6% positives and 52.4% negatives) based on data from PROSO II.^[46]^ Solubility was estimated by retrospective analysis of electronic laboratory notebooks. The notebooks were part of a large effort called the Protein Structure Initiative and consider sequences linearly through the following stages: Selected, Cloned, Expressed, Soluble, Purified, Crystallized, HSQC (heteronuclear single quantum coherence), Structure, and deposited in PDB.^[52]^ The peptides were identified as soluble or insoluble by “Comparing the experimental status at two time points, September 2009 and May 2010, we were able to derive a set of insoluble proteins defined as those which were not soluble in September 2009 and still remained in that state 8 months later.” ^[46]^

#### 2.1.3 Non-fouling

Data for predicting resistance to non-specific interactions (non-fouling) is obtained from.^[35]^ positive data contains 3,600 sequences. Negative examples are based on 13,585 sequences (20.9% positives and 79.1% negatives) coming from insoluble and hemolytic peptides, as well as, the scrambled positives. The scrambled negatives are generated with lengths sampled from the same length range as their corresponding positive set, and residues sampled from the frequency distribution of the soluble data set. Samples are weighted to account for the class imbalance caused by the dataset size for negative examples. A non-fouling peptide (positive example) is defined using the mechanism proposed in White et al.^[53]^. Briefly, White et al. showed that the exterior surfaces of proteins have a significantly different frequency of amino acids and this increases in aggregation prone environments, like the cytoplasm. Synthesizing self-assembling peptides that follow this amino acid distribution and coating surfaces with the peptides creates non-fouling surfaces. This pattern was also found inside chaperone proteins, another area where resistance to non-specific interactions is important.^[54]^

### 2.2 Model Architecture

To identify the position-invariant patterns in the peptide sequences, we build a recurrent neural network (RNN), using a sequential model from Keras framework^[55]^ and the TensorFlow deep learning library back-end.^[56]^ In specific, the RNN employs bidirectional Long Short Term Memory (LSTM) networks to capture long-range sequence correlations. Compared to the conventional RNNs, LSTM networks with gate control units (input gate, forget gate, and output gate) can learn dependency information between distant residues within peptide sequences more effectively.^[57–59]^ They can also partly overcome the problem of vanishing or exploding gradients in the back-propagation phase of training conventional RNNs.^[60]^ We use a bidirectional LSTM (bi-LSTM) to enhance the capability of our model in learning bidirectional dependence between N-terminal and C-terminal amino acid residues. An overview of the RNN architecture is shown in Figure 2.

**Figure 1:**
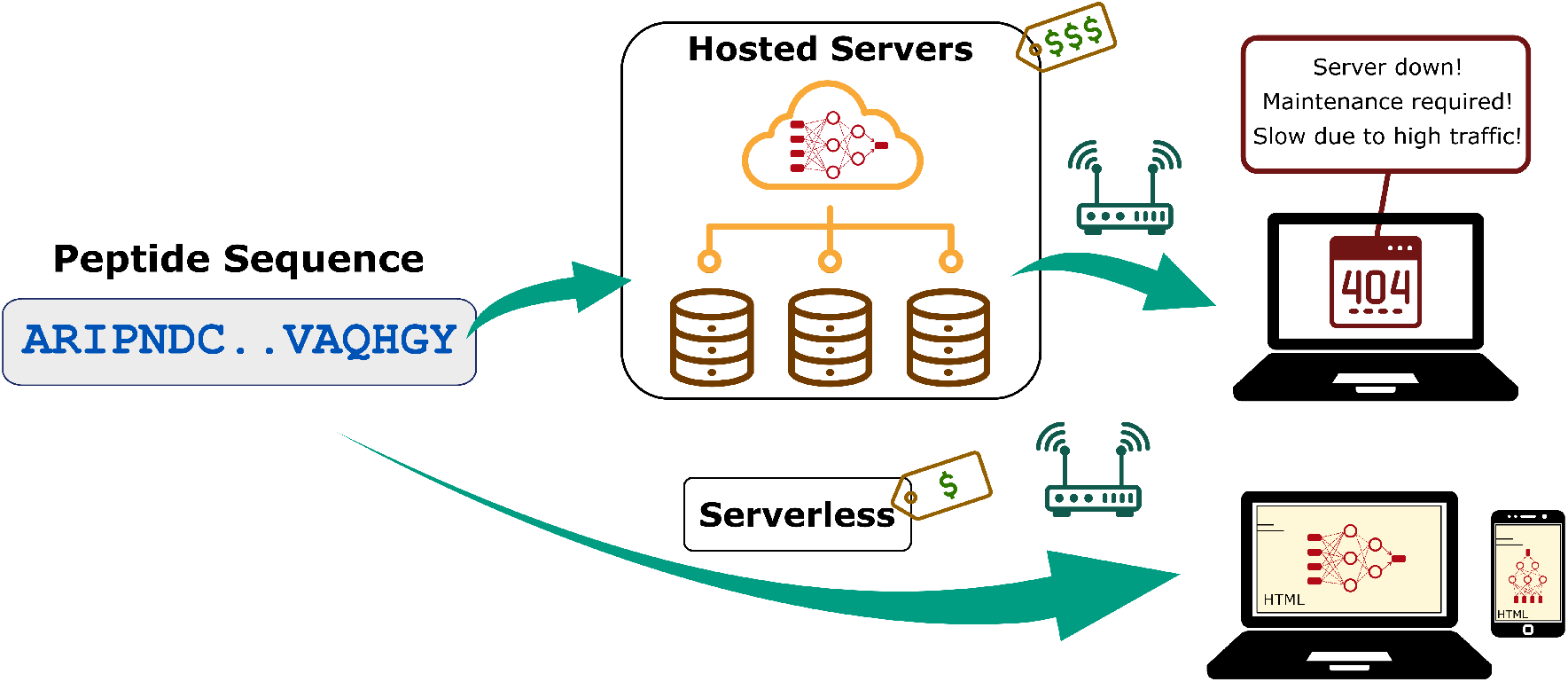
Conventional web-based cheminformatics frameworks vs the proposed serverless approach.

**Figure 2:**
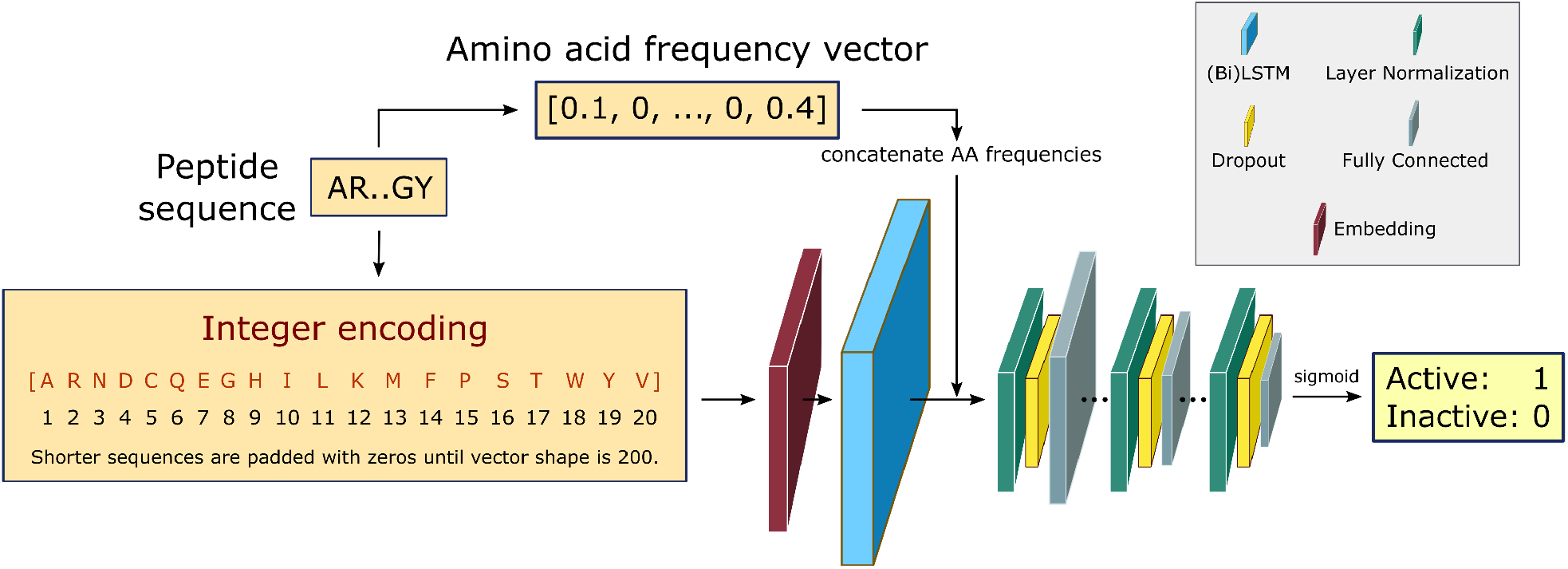
RNN architecture. Fixed-length integer encoded sequences are first fed to a trainable embedding layer, yielding a semantically more compact representation of the input essential amino acids. The bidirectional LSTMS and direct inputs of amino acid frequencies prior to the fully connected layers, improves the learning of bidirectional dependency between distant residues within a sequence. The fully connected layers are down-sized in three consecutive steps with layer normalization and dropout regularization. The final layer uses a sigmoid activation to output a scalar that shows the probability of being active for the desired training task.

Peptide sequences are represented as integer encoded vectors of shape 200, where the integer at each position in the vector corresponds to the index of the amino acid from the alphabet of the 20 essential amino acids: [A, R, N, D, C, Q, E, G, H, I, L, K, M, F, P, S, T, W, Y, V]. Maximum length of the peptide sequence is fixed at 200, and all sequences with higher lengths are excluded. For those sequences with shorter lengths, zeros are padded to the integer encoding representation to keep the shape fixed at 200 for all examples, to allow input sequences with flexible lengths. Note that this is primarily applied to the training step for implementation considerations, and the trained model can make predictions on variable-length sequences as input. Every integer encoded peptide sequence is first fed to an embedding layer. The embedding layer enables us to convert the indices of discrete symbols (i.e. essential amino acids), into a representation of a fixed-length vector of defined size. This is beneficial in the sense of creating a more compact representation of the input symbols, as well as yielding semantically similar symbols close to one another in the vector space. This embedding layer is trainable, and its weights can be updated during training along with the others layers of the RNN.

The output from the embedding layer either goes to a double stacked bi-LSTM layer or a single LSTM layer, to identify patterns along a sequence that can be separated by large gaps. The former is used in predicting solubility and hemolysis, whereas the latter is for predicting peptide’s resistance to non-specific interactions (non-fouling). The rationale behind this choice for the non-fouling model is that the bi-LSTM layer did not contribute to a better performance, when compared with the LSTM layer (same ACC and AUROC of 82% and 0.93, respectively). The output from the LSTM layer is then concatenated with the relative frequency of each amino acid in the input sequences. This choice is partially based on our earlier work,^[61]^ and helps with improving model performance. The concatenated output is then normalized and fed to a dropout layer with a rate of 10%, followed by a dense neural network with ReLU activation function. This is repeated three times, and the final single-node dense layer uses a sigmoid activation function to force the final prediction as a value between 0 and 1. This scalar output shows the probability of the label being positive for the corresponding predicted peptide biological activity. We use this probability to evaluate the confidence of the model in making inference on new sequences in our web-based implementation.

The hyperparameters are chosen based on a random search that resulted the best model performance in terms of the Area Under the Receiver Operating Characteristic (AUROC) curve^[62]^ and accuracy. The AUROC shows the model’s ability to discriminate between positive and negative examples as the discrimination threshold is varied, and the accuracy is defined as the ratio of correct predictions to the total number of predictions made by the model. The embedding layer has the same input dimension of 21 (alphabet length added by one to account for the padded zeros), and output dimension of 32. The LSTM layer has 64 units, and the first, second and third dense layers have 64, 16 and 1 units, respectively. We train with Adam optimizer^[63]^ of binary cross-entropy loss function, which is defined as

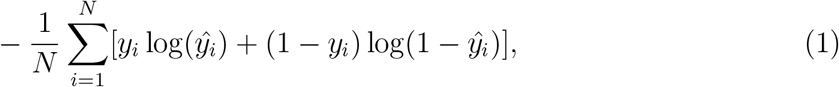

where *y*_*i*_ is the true value of the *i*th example, *ŷ*_*i*_ is the corresponding prediction, and *N* is the size of the dataset. The learning rate is adapted using a cosine decay schedule with an initial learning rate of 10^*−*3^, decay steps of 50 and minimum of 10^*−*6^. Data split for training, validation and test is 81%, 9% and 10%, respectively. To avoid overfitting, we add early stopping with patience of 5 that restores model weights from the epoch with the maximum AUROC on the validation set during training.

Previous models for peptide prediction tasks use a variety of deep learning and classical machine learning methods. The prediction server PROSO II employs a two-layered structure, where the output of a primary Parzen^[64]^ window model for sequence similarity and a logistic regression classifier of amino acid k-mer composition, are fed to a second-level logistic regression classifier. HAPPENN uses normalized features selected by SVM and ensemble of Random Forests, which are fed to a deep neural network with batch normalization and dropout regularization to prevent overfitting. DSResSol (1), that takes advantage of the integration of Squeeze-and-Excitation (SE)^[65]^ residual networks^[66]^ with dilated convolutional neural networks.^[67]^ In specific, the model includes five architectural units, including a single embedding layer, nine parallel initial CNNs with different filter sizes, nine parallel SE-ResNet blocks, three parallel CNNs, and fully connected layers.

## 3 Results

Table 1 shows the classification performance for all the three tasks, along with a comparison between our RNN model and the state-of-the-art methods (see Section 1.1 for a brief overview). All models achieve the same result range as the state-of-the-art methods. We compare the feature extraction capability of our RNN with other unconditional protein language models that provide a pre-trained sequence representations, that transfer well to supervised tasks. In specific, we train two machine learning models on the hemolytic dataset, using UniRep ^[68,69]^ representation of the peptide sequences, followed by a logistic regression, and a Random Forests^[70]^ classifier. Our RNN architecture slightly outperforms both models in terms of AUROC. The one-hot representation of peptides followed by an RNN results the best hemolysis model in terms of AUROC in.^[71]^ The choice of one-hots requires training features specific to each position though, so we do not expect the model to generalize. In contrast, our model is length-agnostic and will have a relatively smaller generalization error for sequences with lengths it has not observed before. Moreover, this removes the need for having training data at each position for each amino acid.

**Table 1:** Performance comparison on the testing dataset. Best performing method for each task is in bold. Our approach is highlighted with an asterisk.

| Method | Task | ACC(%) | AUROC |
| --- | --- | --- | --- |
| Embedding + Bi-LSTM* | Hemolysis | 84.0 | 0.84 |
| UniRep + Logistic Regression | Hemolysis | 82.0 | 0.81 |
| UniRep + Random Forests | Hemolysis | 84.0 | 0.78 |
| HAPPENN<sup>[47]</sup> | Hemolysis | 85.7 | - |
| HLPpred-Fuse<sup>[50]</sup> | Hemolysis | - | **0.91<sup>a</sup>** |
| one-hots + RNN<sup>[71]</sup> | Hemolysis | 76.0 | 0.87 |
| Embedding + LSTM* | Non-fouling | 82.0 | 0.93 |
| Embedding + Bi-LSTM (MahLooL*) | Solubility | 70.0 | 0.76 |
| PROSO II<sup>[46]</sup> | Solubility | 71.0<sup>b</sup> | 0.78<sup>b</sup> |
| DSResSol (1)<sup>[41]</sup> | Solubility | **75.1<sup>c</sup>** | **0.84<sup>c</sup>** |

<sup>a</sup> AUROC is averaged over two reported datasets excluding <6 amino acid residue peptides.
<sup>b</sup> Based on sequence clustering at 90% identity.
<sup>c</sup> Based on test set.<sup>[72]</sup>

Our predictive model for the solubility task, MahLooL 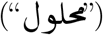, has accuracy of 70.0%, and this is mostly attributed to the difficulty associated with solubility predictions in cheminformatics. Note that the solubility dataset used contains a large distribution of sequence lengths (18-198). DSResSol (1) outperforms all existing solubility models on the testing set from.^[72]^ Readers are encouraged to refer to^[41]^ for the comparison of DSResSol (1) with all the state-of-art sequence-based solubility predictors. For the sake of a better comparison with our approach, we evaluate DSResSol (1)’s performance on the same testing set used for MahLooL. We explore the changes in model performance by training on all the training set, but filtering the testing data based on the sequence lengths, as illustrated in Table 2. MahLooL has comparable performance with respect to DSResSol (1) on the entire testing set. For short length (18-50) peptide, surprisingly it outperforms DSResSol (1), with an AUROC of 0.95, and accuracy of 91.3%. With longer length peptide sequences, the property inference task becomes more difficult by only using the amino acid sequence information, as other experimental settings and conditions become important, adding more epistemic uncertainty to the predictions.

**Table 2:** Performance comparison between MahLooL and DSResSol (1) by training MahLooL on all the training set, while filtering the testing set based on the sequence lengths. Best performing is in bold.

| Length filter | # Test seqs | MahLooL |  | DSResSol (1) |  |
| --- | --- | --- | --- | --- | --- |
|  |  | ACC(%) | AUROC | ACC(%) | AUROC |
| none | 1845 | 70.0 | 0.76 | 71.0 | 0.78 |
| (18-50) | 23 | **91.3** | **0.95** | 87.0 | 0.92 |
| (50-100) | 272 | 76.5 | 0.80 | 75.7 | 0.82 |
| (100-150) | 715 | 70.2 | 0.76 | 70.6 | 0.76 |
| (150-198) | 806 | 70.5 | 0.74 | 71.6 | 0.77 |
| (18-100) | 295 | 78.0 | 0.82 | 76.9 | 0.83 |

To allow for transparency between users and developers, details of the models’ performance, training procedures, intended use and ethical considerations have been incorporated as model cards^[73]^ on https://peptide.bio/. Model cards present information about how the model is trained, its intended use, caveats about its use, and any ethical or practical concerns when using model predictions.

To evaluate the contribution of different architectural components to the model’s performance, we conducted a set of ablative experiments on the solubility model only. In each ablation trial, an architectural component is removed and the corresponding test AUC and accuracy is reported via a 5-fold cross-validation on the solubility dataset. We remove the effect of regularization techniques (see methods in Section 2) in our ablation trials by disregarding the early-stopping callback, and fixing the number of training epochs to 50. The learning rate is also set to a fixed value of 10^*−*3^. This is the reason for the lower performance of the “full model.” The results from our ablation study are shown in Table 4, sorted by the highest AUROC. We point out that the AUROC of the solubility model has a significant drop from 0.76 to 0.68 after removing the regularization callbacks and fixing the learning rate in our cross-validation analysis. Removing amino acid count frequencies, dropout and layer normalization layer both reduced AUROC by about 2%. The removal of the 1^st^ and 2^nd^ dense layers decreased performance by about 5%. Finally, our ablation analysis shows that the Bi-LSTM is the most contributing component of the architecture, as its removal decreased AUROC by about 10%. Indeed, the bidirectionality feature of Bi-LSTM layers boosts the performance by enabling additional learning of the dependence between N-terminal and C-terminal amino acid residues.

**Table 3:** Summary of model cards. Intended use, caveats and any ethical or practical concerns with the three developed models. For more details, refer to https://peptide.bio/.

|  | Hemolysis | Solubility | Non-fouling |
| --- | --- | --- | --- |
| **Intended Use** | Peptides between 1 and 190 residues. L- and canonical amino acids. | Peptides or proteins expressed in E. coli that are less than 200 residues long. May provide solubility predictions more broadly applicable. | Short peptides between 2 and 20 residues. |
| **Factors** | Dataset was from sequences thought to be antimicrobial or clinically relevant. | Solubility was defined in PROSO II<sup>[46]</sup> as sequence that was transfectable, expressible, secretable, separable, and soluble in E. coli system. | The dataset was gathered based on the mechanism proposed in.<sup>[53]</sup> |
| **Ethical Considerations** | These predictions are not a substitute for laboratory experiments. | None noted. | None noted. |
| **Caveats** | The sequences tested were typically from biological sources. | This data is mostly long sequences and so may not be as applicable to solid-phase synthesized peptides. The model accuracy is low for long sequences. | This data is mostly short sequences. The mechanism is indirect. The negative examples have insoluble peptides over-represented so that the accuracy may be inflated if only comparing soluble peptides. |

**Table 4:** Ablation trials to evaluate the contribution of model’s architectural components in the classification performance on the solubility dataset via 5-fold cross-validation. For comparison, the performance of the model with full architecture (as shown in Figure 2) is highlighted with an asterisk.

| Change in Architecture | ACC(%) | AUROC |
| --- | --- | --- |
| None*(full model) | $63.1 \pm 4.1$ | $0.683 \pm 0.046$ |
| Removing AA count frequencies | $62.2 \pm 2.9$ | $0.667 \pm 0.027$ |
| Removing dropout | $61.7 \pm 2.4$ | $0.667 \pm 0.030$ |
| Removing layer normalization | $62.2 \pm 3.4$ | $0.661 \pm 0.030$ |
| Removing 1<sup>st</sup> and 2<sup>nd</sup> dense layers | $59.8 \pm 2.0$ | $0.637 \pm 0.021$ |
| Removing bidirectionality of LSTM layer | $56.1 \pm 1.1$ | $0.580 \pm 0.015$ |

## 4 Discussion

We present three sequence-based classifiers to predict hemolysis, solubility, and resistance to non-specific interactions of peptides, and achieve competitive results compared with state-of-the art models. The hemolytic model predicts the ability for a peptide to lyse red blood cells, and is intended to be applied to peptides between 1 and 190 residues, L- and canonical amino acids (AUROC and accuracy of 0.84 and 84.0, respectively). Hemolysis training dataset is from sequences thought to be antimicrobial or clinically relevant, so it may not generalize to all possible peptides. Our solubility model, MahLooL, is trained with data mostly containing long sequences, thus, it may not be as applicable to solid-phase synthesized peptides. MahLooL provides the state-of-art sequence-based solubility predictions for short peptides (<50) with AUROC and accuracy of 0.95 and 91.3%, respectively. However, its accuracy is lower for long peptide sequences (>100). Its intended use is for peptides or proteins expressed in E. coli that are less than 200 residues long, and may provide solubility predictions more broadly applicable. The non-fouling model predicts the ability for a peptide to resist non-specific interactions, and is intended to be applied to short peptides between 2 and 20 residues (AUROC and accuracy of 0.93 and 82.0, respectively). The non-fouling training data mostly contains short sequences, where negative examples have insoluble peptides overrepresented, so the accuracy may be inflated if only comparing soluble peptides.

## 5 Conclusions

Our proposed RNN models allow for automatic extraction of features from peptide sequences, and removes the reliance on domain experts for feature construction. Moreover, these models are implemented in JavaScript, so that they can run on a static website through a browser on users’ phone or desktop. This *serverless* approach removes the conventional dependence of deep learning models in cheminformatics on third-party hosted servers, thus, reduces cost, increases flexibility, accessibility and promotes open science.

## Acknowledgements

Authors thank Mohammad Madani at University of Connecticut for providing support on the implementation of DSResSol (1). Research reported in this work was supported by the National Institute of General Medical Sciences of the National Institutes of Health under award number R35GM137966. We thank the Center for Integrated Research Computing (CIRC) at University of Rochester for providing computational resources and technical support.

## Data and Code Availability

All data and code used to produce results in this study are publicly available in the following GitHub repository: https://github.com/ur-whitelab/peptide-dashboard. The JavaScript implementation of the models is available at https://peptide.bio/.

## Footnotes

1 https://github.com/ur-whitelab/peptide-dashboard/blob/master/examples/Quick_start.ipynb

